# scID: Identification of transcriptionally equivalent cell populations across single cell RNA-seq data using discriminant analysis

**DOI:** 10.1101/470203

**Authors:** Katerina Boufea, Sohan Seth, Nizar N. Batada

**Author notes:** Correspondence should be addressed to N.B.

## Abstract

The power of single cell RNA sequencing (scRNA-seq) stems from its ability to uncover cell type-dependent phenotypes, which rests on the accuracy of cell type identification. However, resolving cell types within and, thus, comparison of scRNA-seq data across conditions is challenging due to technical factors such as sparsity, low number of cells and batch effect. To address these challenges, we developed scID (*S*ingle *C*ell *ID*entification), which uses the Fisher’s Linear Discriminant Analysis-like framework to identify transcriptionally related cell types between scRNA-seq datasets. We demonstrate the accuracy and performance of scID relative to existing methods on several published datasets. By increasing power to identify transcriptionally similar cell types across datasets with batch effect, scID enhances investigator’s ability to integrate and uncover development, disease and perturbation associated changes in scRNA-seq data.

## Introduction

Single cell RNA-sequencing (scRNA-seq) (Hashimshony et al., 2012; Jaitin et al., 2014; Macosko et al., 2015; Picelli et al., 2014; Ramskold et al., 2012; Zheng et al., 2017) is now being routinely used to characterize cell type specific changes in development, disease and perturbations (Regev et al., 2017) which require analysis of cross-data comparison(Butler et al., 2018; Haghverdi et al., 2018; Kiselev et al., 2018). However, there are technical and biological variations between the different datasets that may hamper the joint analysis of these data (Butler et al., 2018; Haghverdi et al., 2018; Kiselev et al., 2019).

Batch correction approaches such as CCA (Butler et al., 2018) and MNN (Haghverdi et al., 2018) have been designed to integrate scRNA-seq datasets. The advantage of these methods is that batch correction allows merging of datasets which provide more power to resolve cell types (Heimberg et al., 2016), particularly those that are rare. In both CCA and MNN, cells that have similar local correlation or neighbourhood structure in the datasets are paired. However, when the data are significantly imbalanced in the number of cells and library size per cell, both of which influences accuracy of clustering, it is unclear how these methods perform. Moreover, depending on the extent of technical bias or extent of overlap in cell types between the data, untangling technical from biological variation between data may not be possible and runs the risk of over-correcting, i.e. the biological variance between the data may be lost.

As deeply characterized, extensively validated and annotated tissue, organ and organism level atlases are increasingly being generated (Regev et al., 2017), it is worthwhile to reuse such high-quality information to identify known populations of cells in the target data with lower quality. In addition to the propagation of cell labels and metadata across datasets, reference data can also be used to group cells in the target data in which unsupervised clustering does not resolve consistently given the partial knowledge of the cell types in the literature or with the clusters found in the superior quality reference data. In such cases, a biomarker-based approach for identifying transcriptionally equivalent cells across datasets involves determining whether the top marker genes derived from the reference clusters are present in the target cells. However, this biomarker-based approach does not work effectively due to the dropout problem with scRNA-seq data i.e. for a large proportion of genes, the expression values are not sampled (Bacher and Kendziorski, 2016; Stegle et al., 2015; Vallejos et al., 2017). To map cells from target scRNA-seq data to a reference data, scmap (Kiselev et al., 2018) extracts features from each reference cluster and uses a distance and correlation based metric to quantify the closeness between each reference cluster’s centroid and the cell in the target data to assign target cells to reference clusters. However, distance metrics in high dimensional space may not work as expected (Aggarwal, 2001). CaSTLe uses genes with high expression in both the target and reference data or those that have high mutual information to clusters identities in the reference datasets to identify equivalent populations (Lieberman et al., 2018). However, CaSTLe’s accuracy has not been systematically assessed against newer developed methods.

Here we present a novel method called scID for identifying transcriptionally related groups of cells across datasets by exploiting information from both the reference and the target datasets without making any assumptions regarding the nature of technical and biological variation in these datasets. More specifically, scID extracts a list of cluster-specific genes (markers) from the reference dataset, and weighs their relevance in the target dataset by learning a classifier with respect to a putative population of cells either expressing or not expressing those genes. Through extensive analysis of published datasets with batch effect and strong asymmetry in cell number and library sizes per cell in which the independent clustering of the target data via unsupervised method is not obviously similar to that in the reference, we show that scID is more accurate than the above-mentioned methods. Thus, scID helps uncover hidden biological variation present between scRNA-seq datasets that can vary broadly in batch effect and quality (i.e. different number of cells, dropout levels and dynamic range of gene expression).

## Results

### Identification of transcriptionally similar cells across scRNA-seq datasets

Identifying target cells ***g**^j^′s* that match a reference cluster (*C*) can be seen as a binary classification problem *f^C^*(***g**^j^*) by treating the reference cluster-of-interest as one class (*C*) and the rest of the reference clusters as another class (*C*^−^), i.e., *f^C^*(***g**^j^*) ∈ {*C, C*^−^}. Assuming that cell types can be linearly separable, one can, for example, use Fisher’s linear discriminant analysis to learn a direction, 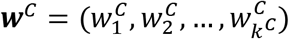, from the reference data that maximises the separation between *C* and *C*^−^ population where *k^C^* is the number of signature genes or features for cluster *C*. Thus, to each target cell we can assign an ‘scID score’

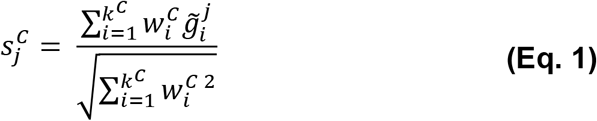

where 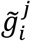 is the normalized gene expression of feature *i* in cell *j* and 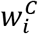 is the weight that reflects the discriminatory power of the *i*-th gene, such that in the projected space, the *C* and the *C*^−^ populations are maximally separated. To mitigate the influence of batch effect, it may be more suitable to estimate weights from the target data instead of the reference. Given putative *c* and *c*^−^ populations selected from the target data, the weights are estimated as

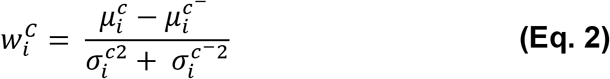

where 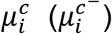 represents mean expression of the putative target cluster *c*(*c*^−^) and 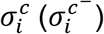 represents standard deviation of gene expression within the putative target cluster *c*(*c*^−^). A target cell can then be assigned to the reference cluster *C* by choosing a suitable threshold. scID maps cells in the target data that are equivalent to clusters in the reference data in three key stages (**Figure 1A**).

**Figure 1.**
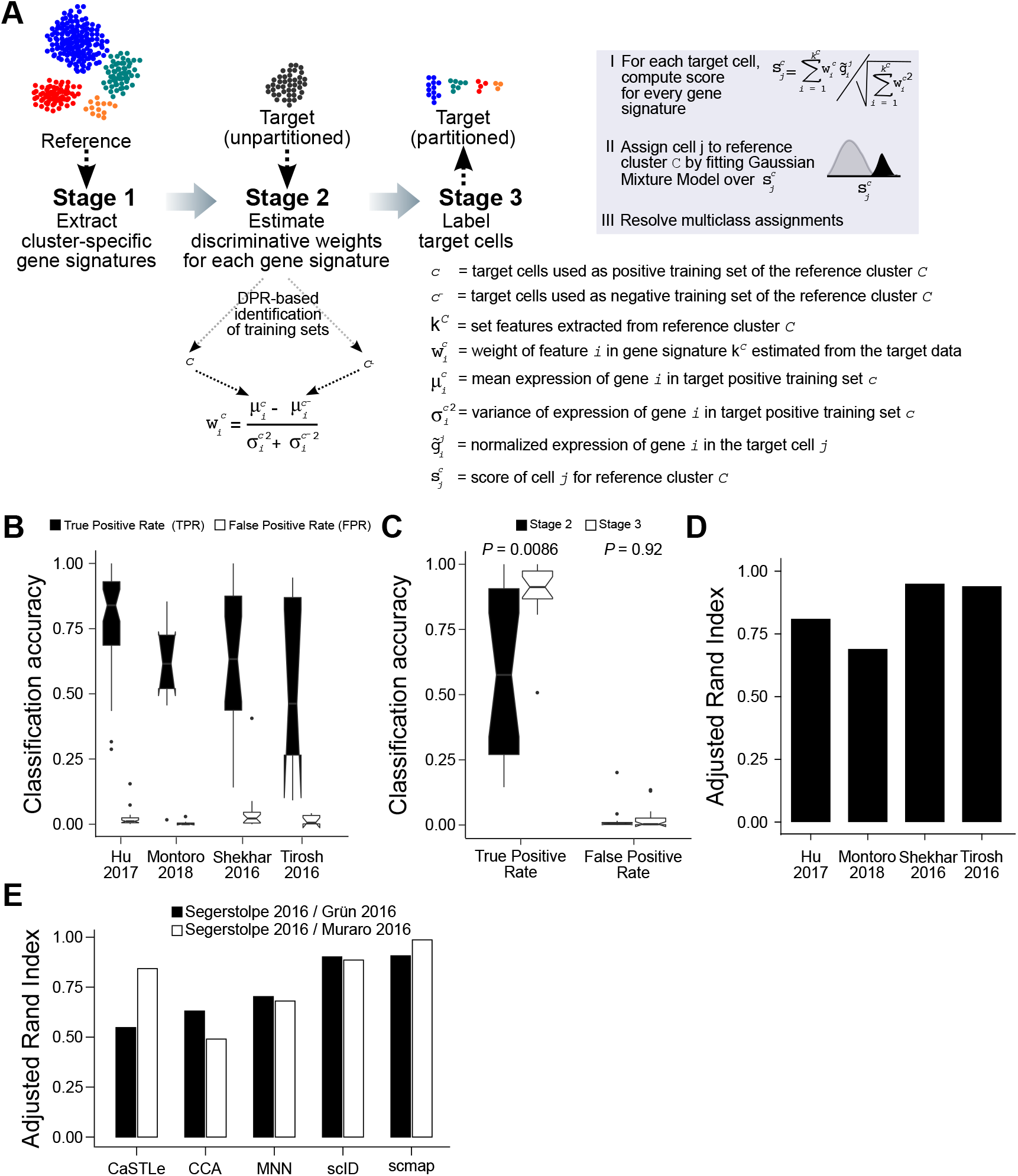
Overview and assessment of scID. **(A)** The three main stages involved in mapping cells across scRNA-seq data with scID are as follows: In stage 1, gene signatures are extracted from the reference data (shown as clustered groups on a reduced dimension). In stage 2, discriminative weights are estimated from the target data for each reference cluster-specific gene signature. In stage 3, every target cell is scored for each feature and is assigned to the corresponding reference cluster. **(B)** Quantification of accuracy of DPR classification (Stage 2 of scID). Boxplot shows interquartile range for TPR (black) or FPR (white) for all the cell types in each published dataset listed in the x-axis. **(C)** Quantification of TPR and FPR of stage 2 (black) and stage 3 (white) of scID. Significance was computed using two-sided paired Kruskal-Wallis test for difference in TPR or FPR between stage 2 and stage 3. **(D)** Assessment of accuracy of scID via self-mapping of published datasets. The indicated published data (x-axis labels) were self-mapped, i.e. used as both reference and target, by scID and the assigned labels were compared to the published cell labels. **(E)** Assessment of classification accuracy of scRNA-seq data integration. Human pancreas Smart-seq2 data (Segerstolpe et al) was used as reference and CEL-seq1 as target (white; Grun et al) or CEL-seq2 as target (black; Muraro et al).

In the first stage, genes that are differentially expressed in each cluster (herein referred to as gene signatures) are extracted from each cluster of the reference data. In the second stage, for each reference cluster *C*, the weights of the genes in the respective gene signature (which reflect their discriminability) are estimated either from the reference or from the target data, depending on the nature of the data, by identifying putative *c* and *c*^−^ cells using an approach we term differential precision-recall (DPR). In the last stage, the extracted features (i.e. the combination of gene signatures and their weights) are used to identify target cells that are most concordant with extracted features. A finite mixture of Gaussians is used to model the scores of the target cells. Cells with high likelihood of falling under the distribution with the highest average score are assigned to the reference cluster from which the gene signatures were extracted. Ideally a target cell should be assigned to only one reference cluster but when closely related cells are present in the reference data, it is possible that the features extracted are not able to highly discriminate between them leading to assignment of a cell to multiple clusters in the reference. Conflicts in assignments are resolved using a relative normalized score strategy. Further details are provided below and in the **Methods** section.

As the accuracy of weights depends strongly on selection of true cells, we assessed the accuracy with which the DPR approach identified true cells in several published reference data. We observed that the median true positive rate of the DPR approach was between 45% and 80% and the median false positive rate was between 0 and 5% (**Figure 1B**). Moreover, the estimated weights after the DPR approach were consistent with weights from the reference data using published label (**Supplementary Figure 1**). Thus, while the DPR approach is stringent and misses higher than desired levels of true cells, its role in the overall scID methodology is the recovery of true cells with very low false positive rate so that the estimated gene weights are accurate. To assess the accuracy of the scoring approach (Stage 3 of scID) and to quantify its contribution to the improvement in classification accuracy, we used the mouse retinal bipolar DropSeq data (Shekhar et al., 2016) and compared the true positive and false positive rate of cells classified by the DPR approach to the subsequent weight-based classification. We find that, relative to the accuracy of the DPR approach (**Figure 1B**), the accuracy of the weight-based classification of cells has significantly higher true positive rate without increase in the false positive rate (**Figure 1C**). To assess the stability and generalizability of scID across scRNA-seq data across a range of quality and cell type composition, we performed self-mapping (i.e. using same dataset as reference and target) of four published datasets spanning different tissues and technologies. The accuracy of self-mapping, as assessed by Adjusted Rand Index, was relatively high (**Figure 1D**).

We next tested our assumption that different target data with different batch effect leads to different gene weights and quantified its impact on classification accuracy. We selected two reference-target pairs from similar tissue so that the differences are largely due to technical factors rather than biological factors. To this end, we sought to detect equivalent cells across multiple human pancreas datasets from different technologies (Smart-Seq2(Segerstolpe et al., 2016), CEL-Seq1(Baron et al., 2016) and CEL-Seq2(Muraro et al., 2016)) in which the published cell identity labels for each dataset is available. We used the Smart-Seq2 as reference data and CEL-Seq or CEL-Seq2 as target. We confirmed that the two reference/target data pairs (i.e. Smart-Seq2/CEL-Seq1 and Smart-Seq2/CEL-Seq2) had different extent of batch effect (**Supplementary Figure 2**). As expected, the scID estimated weights of the same cell type specific genes calculated from the two target datasets are significantly different, yet the overall accuracy of cell type classification for the two target datasets by scID were similar (**Supplementary Figure 3**). scID was one of the two most accurately and consistently well performing methods (**Figure 1E**).

However, these datasets were relatively balanced in terms of cell number and quality and therefore with mild batch effect. We next assessed these methods in dataset pairs with significant asymmetry in cell number and cell coverage between reference and target datasets in cell number and coverage and have stronger batch effect (Buttner et al., 2019) (**Supplementary Figure 2**).

### Reference based identification of equivalent cells by scID across the mouse retinal bipolar neurons scRNA-seq datasets

In general, a large number of cells despite shallow coverage per cell provide more power to resolve highly similar cells than low number of cells with deep coverage per cell (Heimberg et al., 2016). We sought to identify equivalent cell type in scRNA-seq data strong imbalance in number of cells and coverage per cell (Shekhar et al., 2016) were generated from mouse retinal bipolar neurons using two different technologies: Drop-seq (Macosko et al., 2015) (with ~26,800 cells and ~700 genes per cell) and the plate-based Smart-seq2 (with 288 cells and ~5000 genes per cell). While unsupervised clustering of the full Drop-seq data with Seurat resulted in ~18 discrete clusters, unsupervised clustering of the Smart-seq2 data with Seurat and several other clustering methods (data not shown) identified only 4 clusters (**Figure 2A**) possibly due to low cell numbers.

**Figure 2.**
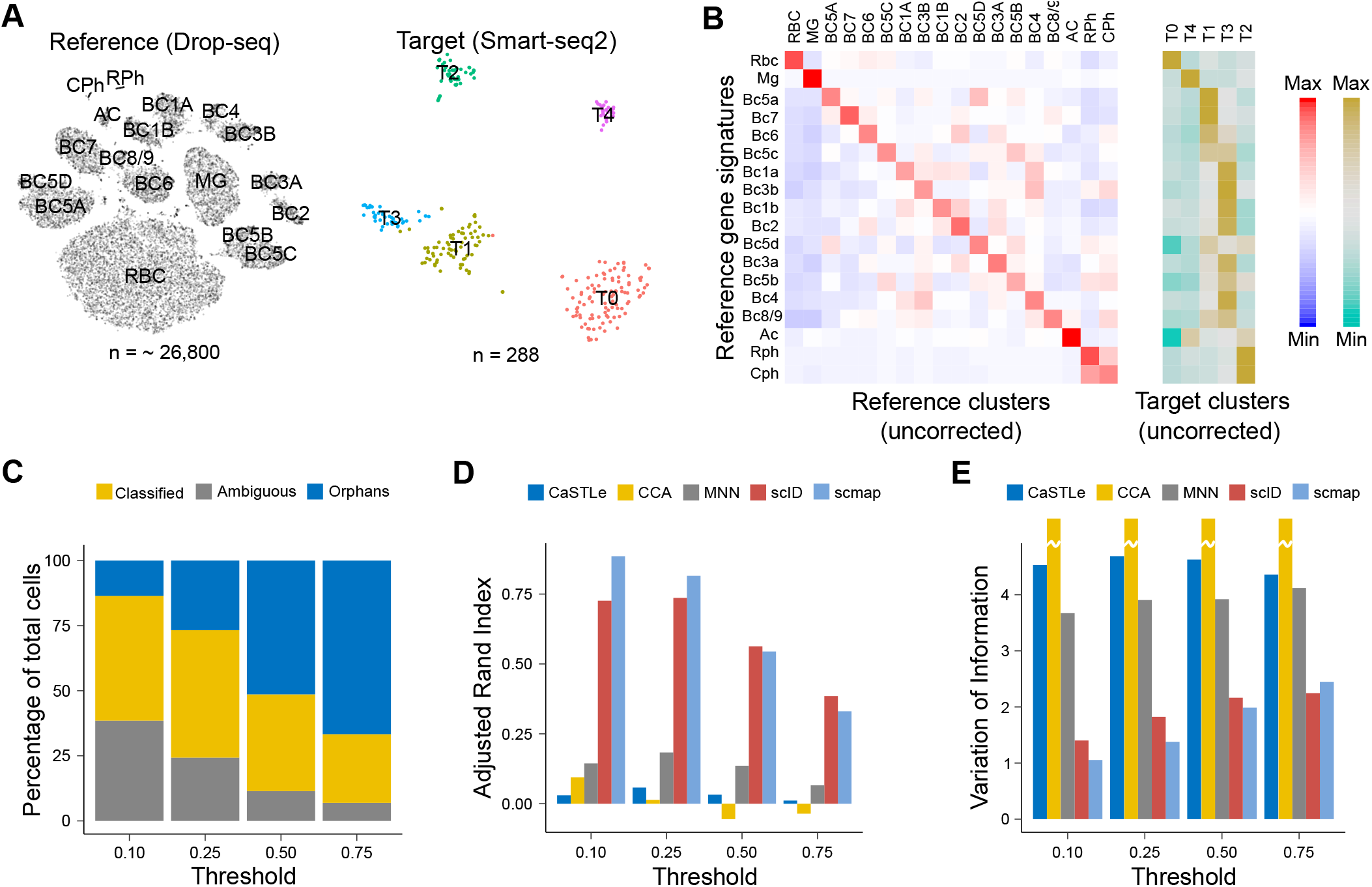
Reference based identification of equivalent cells across the mouse retinal bipolar neurons scRNA-seq datasets. (**A**) t-SNE plot showing clusters in Drop-seq (reference) and Smart-seq2 (target) data of mouse retinal bipolar cells from Shekhar et al. Cluster membership of reference cells (~26,800 cells) were taken from the publication. Smart-seq2 data (288 cells) were clustered using Seurat and clusters name were assigned arbitrarily. (**B**) Heatmap showing z-score normalized average expression of gene signatures (row) in the clusters (column) of the reference Drop-seq data (left) and in the target Smart-seq2 data (right). Red (khakhi) indicates enrichment and blue (turquois) indicates depletion of the reference gene signature levels relative to average expression of gene signatures across all clusters of reference (target) data. (**C**) Identification of target (Smart-seq2) cells that are equivalent to reference (Drop-seq) clusters using marker based approach. The top two differentially enriched (or marker) genes in each reference (Drop-seq) cluster were used to identify equivalent cells in the target (Smart-seq2) data using a thresholding approach. Bars represent percentage of classified and unassigned cells using various thresholds for normalized gene expression of the marker genes as indicated on the x-axis. Gray represents the percentage of cells that express markers of multiple clusters; yellow represents the percentage of cells that can be unambiguously classified to a single cluster; and blue represents the percentage of cells that do not express markers of any of the clusters. These cells are referred to as orphans. X-axis represents different thresholds of normalized gene expression (see **Methods**). (**D, E**) Assessment of accuracy of various methods for classifying target cells. Labels of unambiguously classified cells using marker-based approach (**Figure 2C**, yellow) were used as ground truth. White lines through the bars represent extremely large value.

To determine how the transcriptional signatures of the reference clusters are distributed in the target data, we computed the average gene signature per cluster (**Figure 2B**; see **Methods**). The dominant diagonal pattern in the gene signature matrix for the reference data indicates the specificity of the extracted gene signatures. All the subtypes of bipolar cells (BCs) that were well separated in the reference Drop-seq data co-clustered in the target Smart-seq data (**Figure 2B**). Surprisingly, despite over 7 times greater number of genes per cell in the target Smart-seq2 data compared to the reference Drop-seq data, the biomarker approach of using top markers of clusters from the reference Drop-seq data resulted in unambiguous assignment of only a small proportion of cells (**Figure 2C**).

Thus, we assessed the ability of scID and competing methods to assign cluster identity to cells in the target Smart-seq2 data using Drop-seq data as reference. scID and CaSTLe assigned 100% (n = 288) of the Smart-seq2 cells into 15 Drop-seq clusters while scmap assigned only 63.2% (n=182) of the Smart-seq2 cells into 10 clusters. Using target cells that specifically expressed cell type specific markers (yellow, **Figure 2C**) as ground truth, two independent metrics for accuracy suggested that scID was one of the two most accurately performing methods on this datasets (**Figure 2D,E**).

### Reference based identification of equivalent cells in scRNA-seq and ultra-sparse single nuclei RNA-seq from mouse brain

In many clinical settings where only post-mortem tissues are available, single nuclei-RNA seq (snRNA-seq) (Habib et al., 2017; Lake et al., 2016) may be possible. However, the transcript abundance in the nuclei is significantly lower than in the cytoplasm and as a consequence the complexity of scRNA-seq libraries from nuclei-scRNA-seq from the same tissue is significantly lower than in whole cell scRNA-seq (Habib et al., 2017) thus making it challenging to cluster despite large number of cells. To assess how well scID performs compared to alternatives in this setting, we obtained publicly available Chromium 10X (Zheng et al., 2017) based scRNA-seq data on mouse brain cells and snRNA-seq on mouse brain nuclei (see **Methods**).

The brain whole cells scRNA-seq data had ~9K cells and the brain nuclei snRNA-seq data had ~1K cells. Unsupervised clustering of the data separately identified different number of clusters which did not have an obvious one-to-one correspondence (**Figure 3A**). As the whole cell scRNA-seq data has more cells than the nuclei-seq, we used the former as reference and obtained gene sets that are differentially expressed in each cluster compared to the rest of the cells (**Figure 3B**). Efforts to identify cluster membership of nuclei-seq data based on top markers in each of the clusters from the whole cell scRNA-seq data allowed unambiguous classification of only a small number of cells possibly due to shallow coverage which leads to low dynamic range in gene expression. Not surprisingly, there was significant number of orphan cells (i.e. those in which transcripts of cell markers were not captured) (**Figure 3C**).

**Figure 3.**
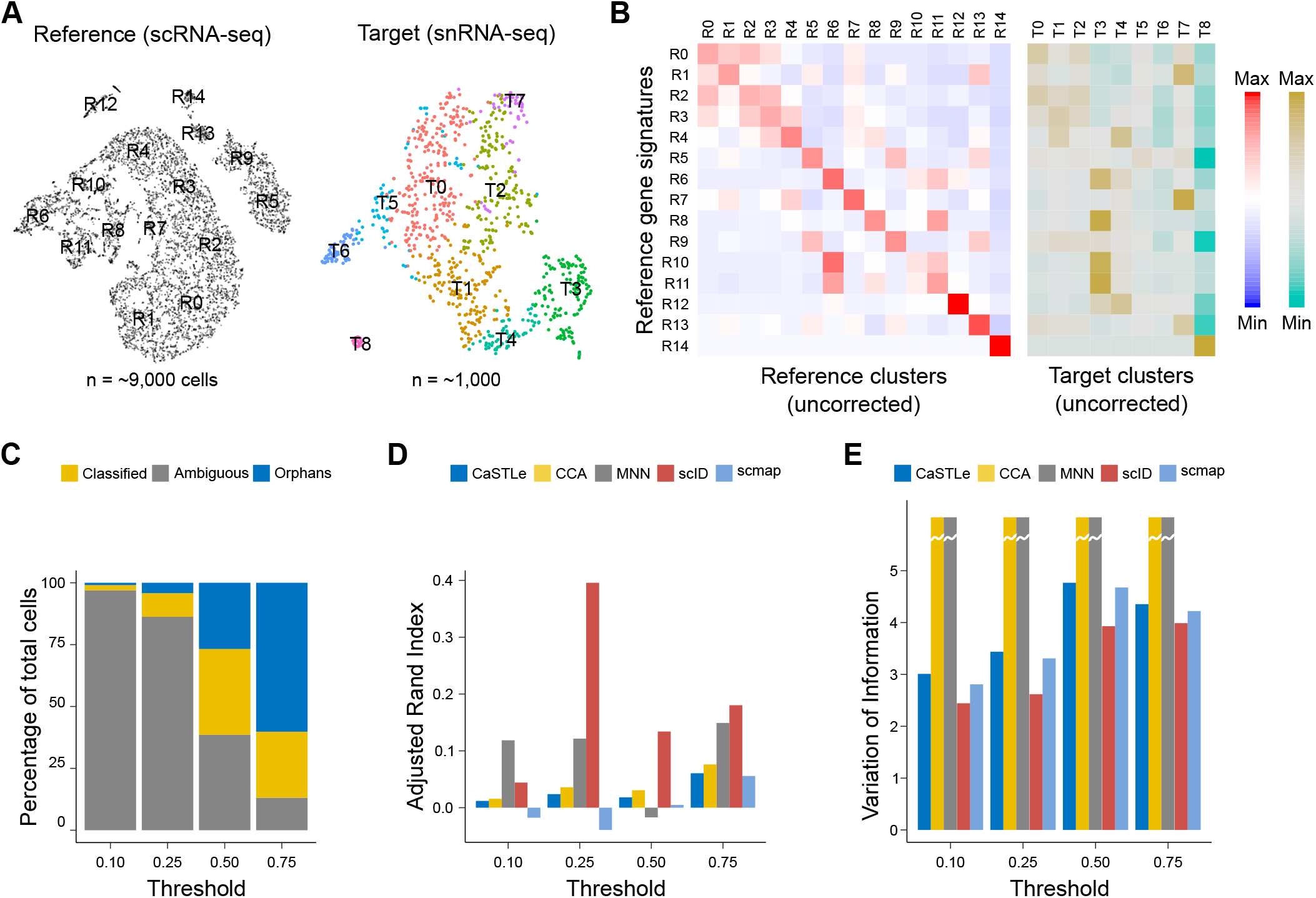
Reference based identification of equivalent cells in single cell and nuclei RNA-seq from mouse brain. (**A**) t-SNE plot showing clusters in the mouse brain scRNA-seq data with ~9,000 cells (left) and tSNE of the mouse brain single nuclei RNA-seq (snRNA-seq) data with ~1,000 cells (right). Data were clustered with Seurat (v3). (**B**) Heatmap showing z-score normalized average expression of gene signatures (rows) in the reference (left) and in the target (right) clusters (columns). Red (khakhi) indicates enrichment and blue (turquois) indicates depletion of the gene signature levels relative to average expression of that gene signature across all clusters of reference (target) data. (**C**) Marker-based identification of cell types in target data. Data was binarized using different thresholds (see methods) that represent expression value of each marker gene relative to the maximum. Two most differentially expressed markers from each reference cluster were used. Cells that express markers of multiple clusters (gray) are labelled as Ambiguous. Cells that only express markers of a single cluster (yellow) are labelled as Classified. Cells that do not express markers of any clusters (blue) are labelled as Orphans. (**D, E**) Assessment of accuracy of mapping methods. Target data that can be unambiguously labelled using marker based approach as shown in **Figure 3C** (yellow) was used as ground truth. To be unbiased, several thresholds of gene expression values (normalized to the maximum expression of that gene) are used as indicated in the x-axis. Higher Adjusted Rand Index and lower Variation of Information indicate better accuracy.

Thus, we assessed the ability of scID and competing methods to assign cluster identity to cells in the target (snRNA-seq) data using scRNA-seq data as reference. scID assigned 99.5% (*n* = 949) of the snRNA-seq cells into 14 scRNA-seq clusters while scmap assigned 60.2% (n = 574) of the snRNA-seq cells into 11 clusters and CaSTLe assigned 100% (n = 954) of the snRNA-seq cells into 15 clusters. Using target cells that specifically expressed cell type specific markers (yellow, **Figure 2C**) as ground truth, two independent metrics for accuracy suggested that scID had the highest accuracy (**Figure 3D,E**).

## Discussion

Comparison of multiple scRNA-seq datasets across different tissues, individuals and perturbations is necessary in order to reveal potential biological mechanisms underlying phenotypic diversity. However, comparison of scRNA-seq datasets is challenging because even scRNA-seq data from the same tissue but with different technology or donor can have significant batch effect confounding the technical with the biological variability. scRNA-seq datasets generated from different scRNA-seq methods and quality (i.e. similar number of cells and library depth per cell) are in general much easier to correct than data generated from different methods and quality. Particularly, in cases of strong asymmetry in quality between datasets to be compared, mapping target cells across data at the individual cell level rather than performing batch correction can be a more appropriate route for identifying known transcriptionally related cells.

We have developed scID which uses the framework of Fisher’s linear discriminant analysis to map cells from a target data to a reference dataset (**Figure 1A**). scID adapts to characteristics of the target data in order to learn the discriminative power of cluster-specific genes extracted from the reference in the target data. scID down-weighs the genes within the signature that are not discriminatory in the target data. Through extensive characterization of multiple published scRNA-seq datasets, we assessed scID’s accuracy relative to alternative approaches in situations where the reference and target datasets have strong asymmetry in quality and have shown that scID outperforms existing methods for recovering transcriptionally related cell populations from the target data which does not cluster concordantly with the reference.

scID can either estimate gene signature weights from the reference or the target data. When the reference and the target data have strong batch effect, estimation of weights from the target appears to be more accurate. However, in cases where there are very similar cell types in the datasets to be compared, estimation of weights from the reference may to be more suitable. Moreover, if the target data contains a cell type that is only weakly related to but not exactly the cell type in the reference, scID is susceptible to falsely classifying such cells.

Due to reliance on gene signatures to discriminate between clusters, a limitation of scID is that it only works for reference clusters that have distinct gene signatures. For a data set to serve as a reference, we expect well defined clusters by cell type such that cluster specific gene signatures exist and they are mutually exclusive as seen in **Figure 2B** and **Figure 3B**. If neither of the two data are well clustered and contain good gene signatures, scID is not a recommended approach for identification of equivalent cell types.

As the scale of single cell RNA-seq data increases and the numbers of clusters obtained are so large that manual annotation is cumbersome, scID can enable automatic propagation of annotations and metadata across clusters in these datasets. In addition, scID can be used for order cells based on an arbitrary user specified gene list (such as those defining a pathway) without the need for providing information on gene expression levels thus aiding in the interpretation of subset of cells across cell types.

## Author contributions

K.B. and N.N.B. conceived the study. K.B, S.S. and N.N.B. developed the computational framework. K.B. implemented the method and performed the analysis. K.B., S.S. and N.N.B. interpreted the results. N.N.B. wrote the manuscript with input from K.B and S.S. N.N.B. supervised the study.

## Acknowledgements

K.B. and N.N.B. were funded by the Chancellor’s Fellowship from the University of Edinburgh to N.N.B.

## Methods

### scID workflow

Given a set of 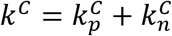 features (genes) that are positive 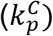 and negative 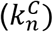 markers for a reference cluster *C*, scID selects target cells equivalent to reference cluster *C* based on their score 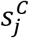, which is a weighted linear sum given by

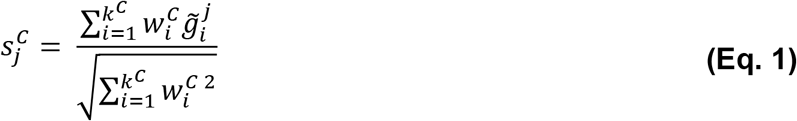

for *j* = 1,…,*n* where *n* is the number cells in the target data, 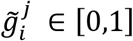 is the normalized gene expression value of the *i*-th gene in the *j*-th cell, and 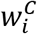 is a weighting factor that represents the discriminative power of gene *i* to identify target cells equivalent to reference cluster *C*. To reduce sensitivity to outliers, the gene expression values are normalized to [0,1] by the 99^th^ percentile instead of the maximum i.e. 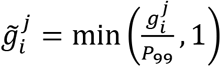, where 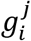 is the library depth normalized gene expression of gene *i* in cell *j* and *P*_99_ is the 99^th^ percentile of the expression of gene *i* across all target cells.

The weights can be computed from the reference data as follows

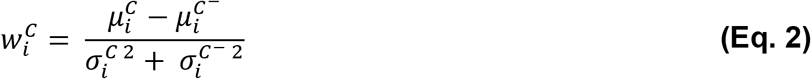

where 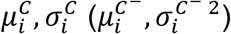 represent the mean and standard deviation, respectively, of gene *i* in the cluster *C* (all clusters except *C*). Each term of the weight of gene *i* is in turn calculated as follows:

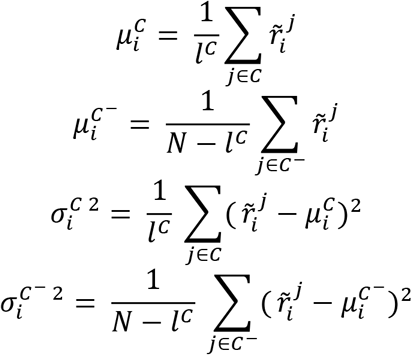

where *l^C^* is the number of cells in cluster *C, N* is the total number of cells in the reference data, and 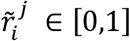 is the normalized gene expression value of the *i*-th gene in the *j*-th cell in the reference data.

This definition of the weights (**Eq. 2**) is a solution to Fisher’s linear discriminant analysis assuming diagonal covariance, where the separation between the two classes is defined as the ratio of the variance between the classes to the variance within the classes. Ideally, we would like to include the full covariance matrix in order to better capture the relationships among clusters. However, we have chosen to approximate the covariance matrix as diagonal for computational efficiency but also due to limitations posed by the nature of scRNA-seq data. When datasets have sparse coverage, the covariance matrix is not full rank and cannot be inverted. Additionally, when the list of genes is long, computing the full covariance matrix is error prone due to insufficient number of samples.

Choosing the weights in this way penalizes high variability and low mean expression in order to account for the following cases:

a. When a positive marker is not expressed uniquely in the *C* population, 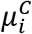 will increase, reducing the weight as it does not provide sufficient evidence of belonging to the cells of interest.
b. When a positive marker is expressed only within a subpopulation of *C*, 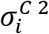 will increase, reducing the weight so that a cell’s score does not drop sharply when the gene is missing. This also accounts for genes with high dropout rate even though they might be specific and sensitive markers.
c. Finally, non-discriminative genes will be down-weighted, as the numerator 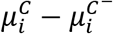 will be low.

Although we can compute these weights from the reference, computing them from the target data can lead to improved accuracy, due to adjustments to the data quality and cell type composition in the target data. However, to do this we need to select target cells that are likely equivalent to the reference cluster *C*. The target cells (denoted as *c*) that express the 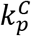 signature genes precisely and specifically and do not express the 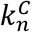 signature genes are selected as equivalent to the reference cluster *C* by clustering the data in the differential precision–differential recall domain which we refer to as differential precision recall or DPR approach.

In general, precision of a cell expressing a set of genes is defined as the total number of expressed genes that belong to the given set divided by the total number of expressed genes and recall is defined as the number of expressed genes that belong to the set divided by the total number of genes in the set.

So, for the set of positive markers (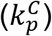

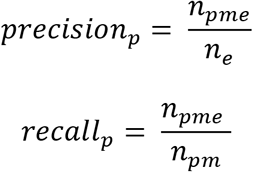

where

*n_pme_*: number of upregulated signature genes expressed

*n_pm_*: total number of upregulated signature genes

*n_e_*: total number of expressed genes

A cell expressing all positive markers will have *recall_p_* = 1 and a cell expressing only positive markers (and no other genes) will have *precision_p_* = 1. Thus, cells equivalent to cluster *C* will be in the first quadrant of the (*precision_p_, recall_p_*) space close to (1,1) and other cells not expressing the positive markers will be close to (0,0).

For the set of negative markers 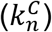

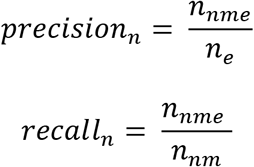

where

*n_nme_*: total number of downregulated signature genes expressed

*n_nm_*: total number of downregulated signature genes

*n_e_*: total number of expressed genes

Similar to precision-recall for positive markers, a cell expressing all negative markers will have *recall_n_* = 1 and a cell expressing only negative markers will have *precision_n_* = 1. Thus, cells equivalent to cluster *C* should be in the third quadrant of the (*precision_n_, recall_n_*) space close to (0,0).

To identify target cells equivalent to reference cluster *C*, we have combined the precision and recall for positive and negative markers via a differential precision (*DP*) and differential recall (*DR*) metric as follows:

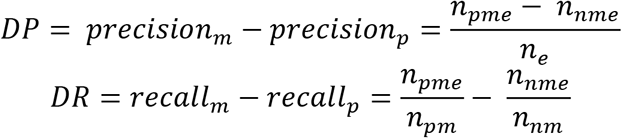

Thus, target cells equivalent to cluster *C* will be close to (1,1), cells very different from cluster *C* will be close to (−1,−1), and cells belonging to clusters that are similar to cluster *C* and share markers will be around (0,0). This will help separate cells equivalent to cluster *C* from cells belonging to very similar clusters that could be otherwise grouped together if we were only using the positive markers.

To select putative positive and negative training populations from the DP-DR space, we cluster the cells using different Gaussian finite mixture models(Scrucca et al., 2016) and select the model with the lowest Bayesian Information Criterion (BIC). The clusters with highest DP and/or DR are selected as candidate matching clusters. From the candidate clusters, we select as *c* the one with the closest to (1,1) centroid as measured by Euclidean distance. All other candidate clusters are discarded from the training set and the remaining clusters are used as training non-matching clusters (*c*^−^).

Analogous to computing the weights in the reference data, they can be computed from the target data using the training sets of cells (*c* and *c*^−^).

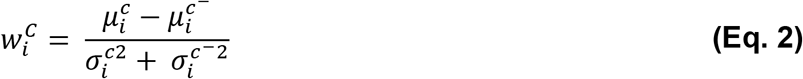

where 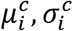 represent the mean and standard deviation, respectively, of gene *i* in the cluster *c* and 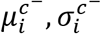 represent the mean and standard deviation, respectively, of gene *i* in the cluster *c*^−^. Each term of the weight of gene *i* is in turn calculated as follows:

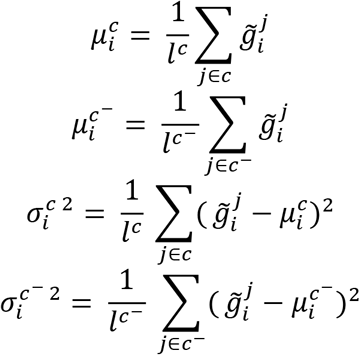

where *l^c^, l*^*c*^−^^ are the number of cells in clusters *c* and *c*^−^ respectively.

The weights estimated from the target data reflect the cellular composition of the target data and quality (i.e. the distribution of dropout and dynamic range of gene expression) of the target data both of which influence the discriminatory power of the features.

Then, to identify target cells equivalent to reference cluster *C*, we fit different finite mixtures of Gaussians on the score 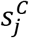 (**Eq. 1**) and assign cells that belong to the population with highest average score to reference cluster *C*.

When the reference data contains clusters that are highly similar transcriptionally, the features from a cluster can be correlated with the features from another cluster which in turn lead scID to assign target cells to multiple reference classes. To resolve this, the scores of target cells 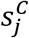 is first z-score normalized and the ambiguous cell is assigned to the reference cluster with the highest normalized score over all other reference clusters it was assigned to.

### Data source

*Human Metastatic Melanoma immune cells from Tirosh et al. 2016* (Tirosh et al., 2016): This Smart-seq2(Picelli et al., 2013) data consists of malignant, immune and stromal cells from metastatic melanoma tumours from 19 patients, a total of 4,645 cells. We have used the 3,254 immune (CD45^+^) cells for our analysis, which on average had 3,925 genes per cell. Data was downloaded from the Broad Institute Single Cell Portal.

*Mouse Retinal Bipolar Neurons from Shekhar et al. 2016* (Shekhar et al., 2016): This study performed Drop-seq and Smart-seq2 experiments on Vsx2-GPF mouse retinal cells. The Drop-seq data had 27,499 cells with an average of 880 genes per cell. The Smart-seq2 data had 288 cells with an average of 4,556 genes per cell. Gene expression data was downloaded from the Broad Institute Single Cell Portal.

*Brain cells from E18 mouse*: This 10X data consists of brain cells from the cortex, hippocampus and subverticular zone of an E18 mouse. The scRNA-seq dataset had 9,128 cells with an average of ~2,500 genes per cell and the single nuclei RNA-seq data have 954 cells with an average of 2,832 genes per cell. Both these datasets were downloaded from 10X Genomics (https://support.10xgenomics.com/single-cell-gene-expression/datasets).

*Murine tracheal epithelium cells from Montoro et al. 2018* (Montoro et al., 2018): This combination of plate-based and droplet-based scRNA-seq data of murine airway epithelial cells consists of 7,193 cells with an average of 1,712 genes per cell. The cells were partitioned into seven clusters annotated post hoc using a biomarker approach by the authors. Data was downloaded from the Broad Institute Single Cell Portal.

*Mouse brain cells from Hu et al. 2017* (Hu et al., 2017): This Drop-seq single nuclei RNA-seq data from cortical tissues of adult mice consists of 18,194 cells with 1,649 genes per cell on average that were partitioned into 40 annotated clusters. Data was downloaded from the Broad Institute Single Cell Portal.

*Unstimulated and stimulated PBMCs from Kang et al. 2018* (Kang et al., 2018): This 10X data consists of 14039 human PBMCs from eight patients, split into two groups; one control and one stimulated with interferon-beta (IFN-β). Seurat CCA was used to align and cluster the data in order to obtain gold standard cell identities as shown in (Butler et al., 2018).

*Human pancreatic islet cells from Segerstolpe et al. 2016* (Segerstolpe et al., 2016): This Smart-Seq2 data consists of pancreatic tissue and islets from six healthy individuals and four type 2 diabetes patients. RPKM-normalized gene expression data and cell labels were downloaded from ArrayExpress (E-MTAB-5061).

*Human pancreatic islet cells from Grün et al. 2016* (Grun et al., 2016): This CEL-seq data consists of pancreatic cells from deceased organ donors with and without type 2 diabetes. Gene expression data and cell labels were downloaded from NCBI GEO (GSE81076).

*Human pancreatic islet cells from Muraro et al. 2016* (Muraro et al., 2016): This CEL-seq2 data consists of islets from cadaveric pancreas. Gene expression data and cell labels were downloaded from NCBI GEO (GSE85241).

### Data Normalization

When datasets were obtained as UMI counts, Counts Per Million (CPM) library-depth normalization was performed prior to the analysis. For the biomarker-based approach we further normalized the gene expression to [0,1] by the 99^th^ percentile in order to use the same threshold between marker genes that can differ significantly in their expression level.

### scID implementation

The R implementation and tutorial for scID is available on Github (https://batadalab.github.io/scID/).

scID has the following user-specified options:

1. `logFC`: Log-fold-change threshold for extracting cluster-specific genesets from the reference data. The `logFC` used for extracting gene signatures for the reference datasets used in the figures are as follows: For Hu et al. 2017 (Hu et al., 2017) `logFC` was set to 0.3; for Montoro et al. 2018 mouse tracheal epithelium data (Montoro et al., 2018) `logFC` was set to 0.5; for Shekhar et al. 2016(Segerstolpe et al., 2016) `logFC` was set to 0.7; for Tirosh et al. 2016 metastatic melanoma Smart-seq2 data (Tirosh et al., 2016) `logFC` was set to 0.5; for Segerstolpe et al. 2016 Smart-seq2 human pancreas data (Segerstolpe et al., 2016) `logFC` was set to 0.5; for 10X E18 mammalian brain data (https://support.10xgenomics.com/single-cell-gene-expression/datasets/3.0.0/neuron_1k_v2) `logFC` was set to 0.6.
2. `estimate_weights_from_target`: Estimate weights using the target data by selecting training sets using the precision-recall-like approach. For all figures in the paper we set this to `TRUE`.
3. `only_pos`: Select only upregulated genes from each reference cluster. For all figures in the paper we set this to `FALSE` in order to select both upregulated and downregulated genes.

### Software tools and parameter settings

We used the following R packages and parameters:

#### Seurat (Butler et al., 2018) (version 3.0.1)

We used UMI count data when available and followed the standard Seurat workflow for clustering and data integration with default settings. When normalized data was provided instead, we skipped the `NormalizeData()` function of the workflow.

#### scran (version 1.10.2)

For running MNN (Haghverdi et al., 2018), we used log-transformed CPM-normalized gene expression values. Aligned and merged expression matrixes were clustered using Seurat with default parameters.

#### scmap (Kiselev et al., 2018) (version 1.4.1)

We used log-transformed CPM-normalized gene expression values for both reference and target data. We selected number of highly variable to genes to be used so that the maximum possible number of target cells can be classified, as shown in the following table. Specifically, for the data pairs of Figure 1E and Figure 3 we used 500 highly variable genes and for the data pairs of Figure 2 we used 150.

#### CaSTLe(Lieberman et al., 2018)

For running CaSTLe we used the code provided in https://github.com/yuvallb/CaSTLe/blob/master/CaSTLeMultiClass.R with all predefined parameters. We used log-transformed CPM-normalized gene expression values for both reference and target data.

#### Biomarker-based classification of cells

To assign labels to target cells using the biomarker-based approach we first extracted the top two highly enriched markers (referred to as biomarkers) from each reference cluster, sorted by average log fold change. Then binarized the 99^th^-percentile-normalized gene expression data using different thresholds (0.10, 0.25, 0.50 and 0.75) and checked which marker genes are present in each cell. Cells that expressed biomarkers of different cell types were labelled as “ambiguous”. Cells that expressed biomarkers of a single cell type were assigned to the respective cell type and cells that did not express any biomarker were labelled as “orphans”. Only the uniquely classified cells were used to assess the performance of other methods.

#### Quantification of batch effect between pairs of scRNA-seq data

To measure the extent of batch effect between the reference-target pairs of data used in the manuscript, we used kBET (Buttner et al., 2019) (version 0.99.5). High rejection rate indicates poor mixing of the data.

#### Statistical tests

For testing the improvement of scID Stage 3 versus scID Stage 2 (**Figure 1C**), we tested the difference in True Positive (TPR) and False Positive Rate (FPR) between the two stages for each reference cluster using two-sided paired Kruskal-Wallis test.

For the comparison of the various methods classification of cells to their gold standard labels we have used the Adjusted Rand Index (ARI) and the Variation of Information (VI) metrics. High similarity between the testing method and the true classification results in high ARI and low VI values.

#### Reproducibility

All the scripts to generate the figures shown in this manuscript are available at https://github.com/BatadaLab/scID_manuscript_figures.

