## Supplementary Figure 1 for "scID: Identification of transcriptionally equivalent cell populations across single cell RNA-seq data using discriminant analysis"

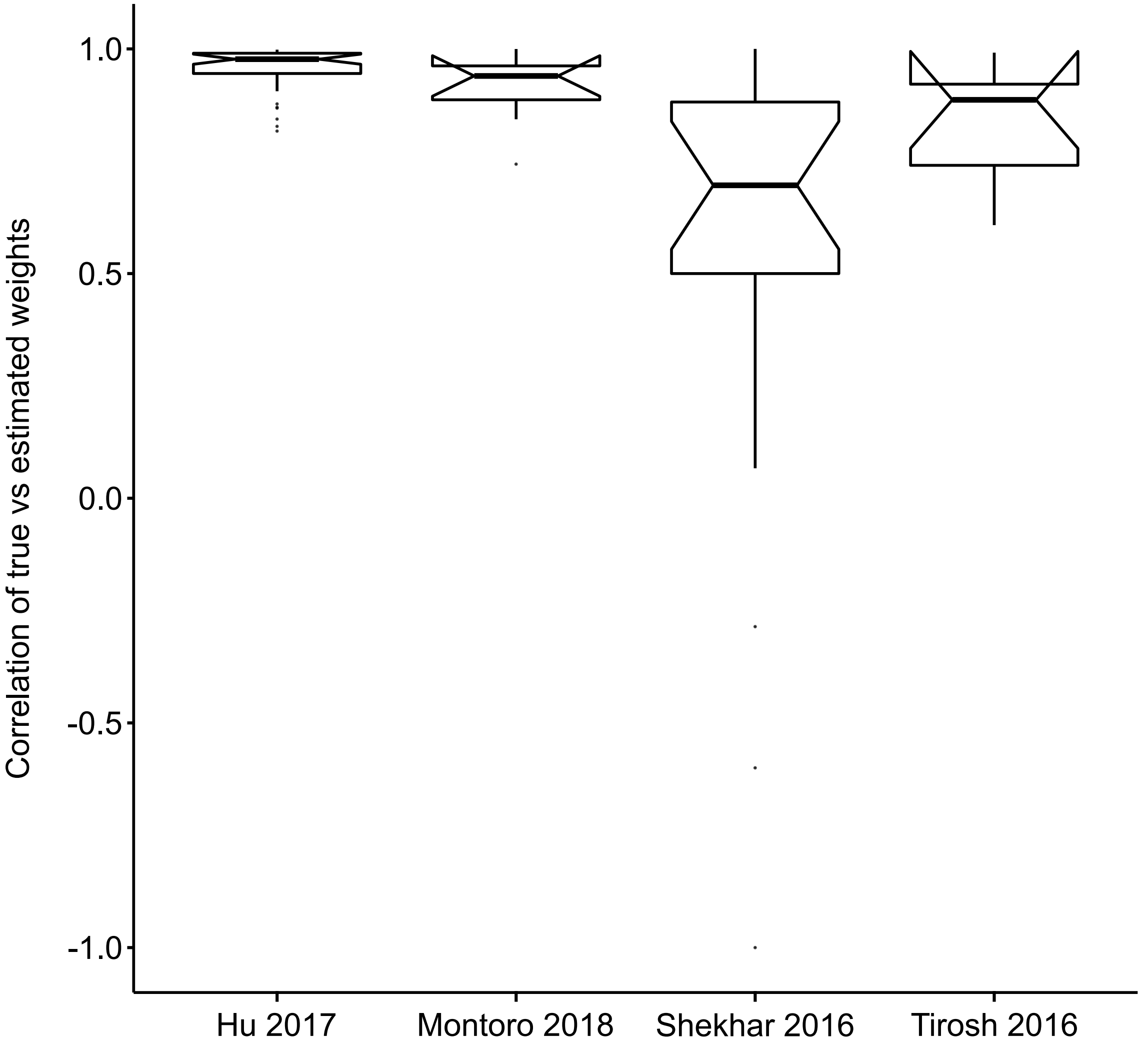


**Supplementary Figure 1. Assessment of accuracy of DPR strategy for cell selection.** Y-axis shows Spearman rank correlation of weights estimated from DPR strategy to weights estimated from the known labels of the reference data. Dataset sources used as both reference and target are listed in the x-axis labels.
