## Supplementary Figure 2 for "scID: Identification of transcriptionally equivalent cell populations across single cell RNA-seq data using discriminant analysis"

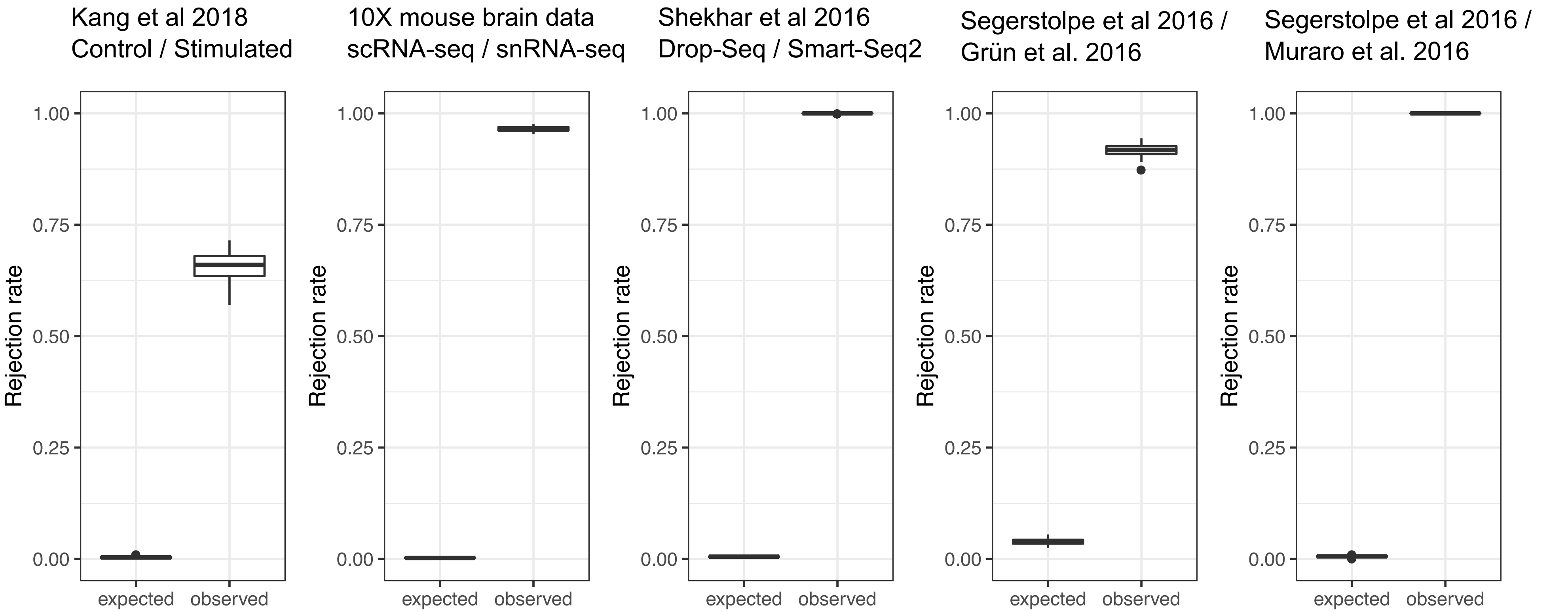


**Supplementary Figure 2. Quantification of batch effect in the reference/target dataset pairs used in the manuscript.** Extent of batch effect as measured by kBET (Büttner et al. 2019, Nature Methods) is shown in the y-axis. The larger the difference between observed and expected rejection rate, the bigger the batch effect. Datasets used are listed in the title of the panels. Kang 2018, which is PBMCs from 8 individuals which were unstimulated and stimulated with interferon-beta and very balanced in terms of cell number and coverage, was used as an example of a dataset with low batch effect
