## Supplementary Figure 3 for "scID: Identification of transcriptionally equivalent cell populations across single cell RNA-seq data using discriminant analysis"

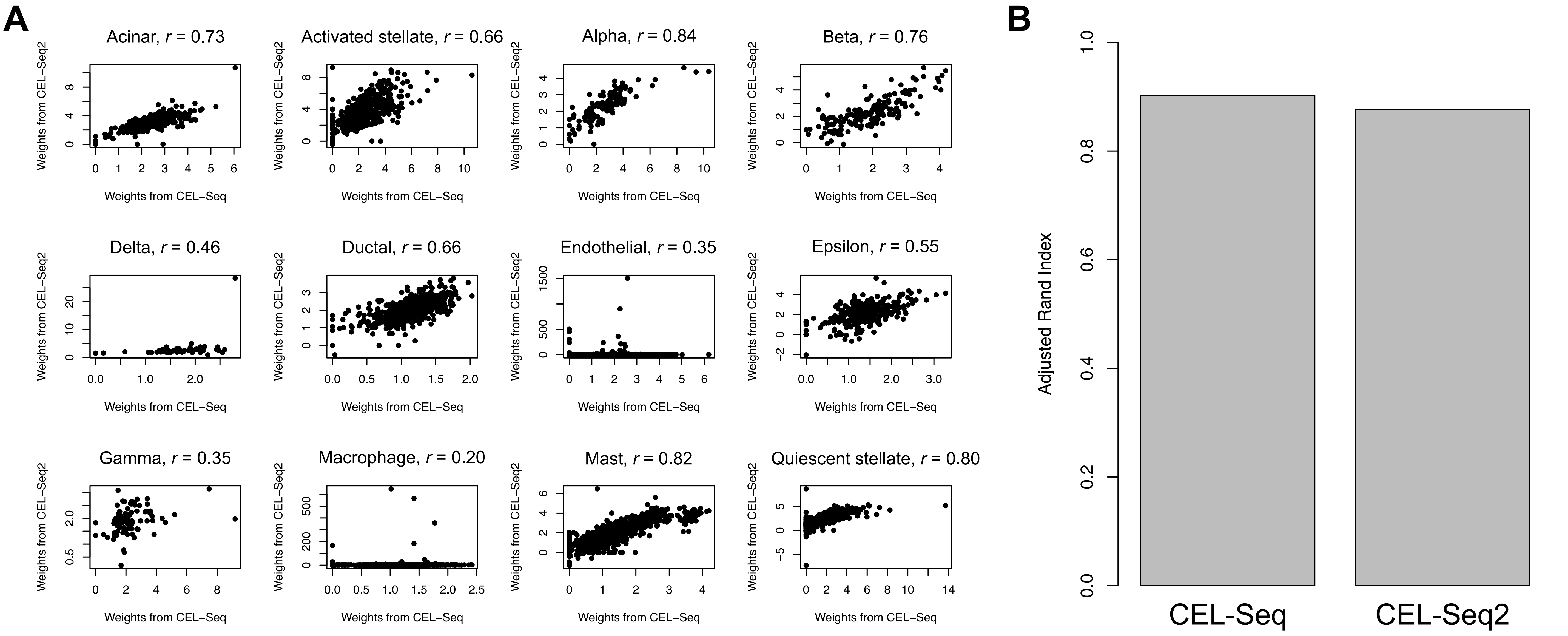


**Supplementary Figure 3. Different target datasets with same cell types have different weights but the classification accuracy of scID is similar. (A)** Scatter of weights estimated from pancreas scRNA-seq CEL-Seq data () is shown on x-axis and estimate from pancreas scRNA-seq CEL-Seq2 data is shown on y-axis using pancread scRNA-seq SMart-Seq2 data (Segerstolpe et al. 2016) as reference. For each of the cell types in the reference data (indictaed in the title of each panel), gene weights were computed using DPR classification in the two target cells. Spearman rank correlation is shown in the title of each pane. Divergence of the correlation from *r* =1 suggests that the weights are not exactly the same in the two target datasets for the same cell type and gene signature.
(**B**) Accuracy of final classification for the two target datasets, calculated using the Adjusted Rand Index.
